# Dissociable default-mode subnetworks subserve childhood attention and cognitive flexibility: evidence from deep learning and stereotaxic electroencephalography

**DOI:** 10.1101/2022.02.25.481973

**Authors:** Nebras M. Warsi, Simeon M. Wong, Jürgen Germann, Alexandre Boutet, Olivia N. Arski, Ryan Anderson, Lauren Erdman, Han Yan, Hrishikesh Suresh, Flavia Venetucci Gouveia, Aaron Loh, Gavin J. B. Elias, Elizabeth Kerr, Mary Lou Smith, Ayako Ochi, Hiroshi Otsubo, Roy Sharma, Puneet Jain, Elizabeth Donner, Andres M. Lozano, O. Carter Snead, George M. Ibrahim

**Affiliations:** Division of Neurosurgery, The Hospital for Sick Children, 555 University Ave., Toronto, Ontario, Canada; Department of Biomedical Engineering, University of Toronto, Toronto, Ontario, Canada; Program in Neuroscience and Mental Health, Hospital for Sick Children, Toronto, Ontario, Canada; Division of Neurosurgery, Toronto Western Hospital, University Health Network, Toronto, Ontario, Canada; Joint Department of Medical Imaging, University of Toronto, Toronto, Ontario, Canada; Maerospace Corporation, Waterloo, Ontario, Canada; Vector Institute for Artificial Intelligence, University Health Network, Toronto, Ontario, Canada; Department of Psychology, The Hospital for Sick Children, 555 University Ave., Toronto, Ontario, Canada, M5G 1X8 Department of Psychology, University of Toronto, Toronto, Canada; Division of Neurology, The Hospital for Sick Children, 555 University Ave., Toronto, Ontario, Canada

## Abstract

**Background:** Cognitive flexibility encompasses the ability to efficiently shift focus and forms a critical component of goal-directed attention. The neural substrates of this process are incompletely understood in part due to difficulties in sampling the involved circuitry.

**Methods:** Stereotactic intracranial recordings that permit direct resolution of local-field potentials from otherwise inaccessible structures were employed to study moment-to-moment attentional activity in children with epilepsy during the performance of an attentional set-shifting task. A combined deep learning and model-agnostic feature explanation approach was used to analyze these data and decode attentionally-relevant neural features. Connectomic profiling of highly predictive attentional nodes was further employed to examine task-related engagement of large-scale functional networks.

**Results:** Through this approach, we show that beta/gamma power within executive control, salience, and default mode networks accurately predicts single-trial attentional performance. Connectomic profiling reveals that key attentional nodes exclusively recruit dorsal default mode subsystems during attentional shifts.

**Conclusions:** The identification of distinct substreams within the default mode system supports a key role for this network in cognitive flexibility and attention in children. Furthermore, convergence of our results onto consistent functional networks despite significant inter-subject variability in electrode implantations supports a broader role for deep learning applied to intracranial electrodes in the study of human attention.

**Funding:** No funds supported this specific investigation. Awards and grants supporting authors include: Canadian Institutes of Health Research (CIHR) Vanier Scholarship (NMW, HY); CIHR Frederick Banting and Charles Best Canada Graduate Scholarship Doctoral Award (SMW); CIHR Canada Graduate Scholarship Master’s Award (ONA); and a CIHR project grant (GMI).

## 1. Introduction

The neural syntax that underlies childhood attention is complex and incompletely understood. Attention relies on the abilities to filter relevant stimuli from the environment (selective attention), maintain focus over time (sustained attention), and to shift focus toward salient stimuli (attentional shift) (Mueller et al., 2017; Newcorn et al., 2001; Smith et al., 2004). Cognitive flexibility refers to an individual’s ability to successfully shift attention in a behaviourally relevant manner (Dajani and Uddin, 2015). Greater cognitive flexibility is associated with improved academic, social, and professional function across the lifespan and its impairment forms a key feature of many neurocognitive and attentional disorders (Dajani and Uddin, 2015; Genet and Siemer, 2011; Roshani et al., 2020; Waltz, 2017).

Shifts in attention are thought to arise from selective synchronization and desynchronization of distinct neural assemblies tuned toward specific behaviours (Womelsdorf and Fries, 2007). Among the distributed networks that subserve attention, a significant body of evidence has supported the role of hierarchical interactions between the salience (SN) and executive control (ECN) networks in guiding attention (Anticevic et al., 2012; Kucyi et al., 2020; Mills et al., 2018). The contribution of the default-mode network (DMN), a critical large-scale functional network that interacts with the SN and ECN, is less clearly defined.

In humans and primates, diverging hypotheses about the role of the DMN have emerged due to observations of both disengagement and recruitment of this network during attentional tasks (Arsenault et al., 2018; Cheng et al., 2020; Crittenden et al., 2015). Andrews-Hanna have proposed a unifying explanation for these findings through functionally segregated subnetworks of the DMN, each with distinct roles in attentional behaviour (Andrews-Hanna et al., 2010).

Indeed, region-of-interest analyses have revealed core DMN subregions comprised of the anterior and posterior cingulate (ACC/PCC) and medial prefrontal cortex (mPFC) that co-activate with attentional shifts in humans (Crittenden et al., 2015; Smith et al., 2019). These functional magnetic resonance imaging (fMRI) findings are however limited by timescale with which granular subnetwork interactions can be resolved (Cohen and Cavanagh, 2011; Combrisson et al., 2017; Pandarinath et al., 2018; Vahid et al., 2020). There is therefore an unmet need to further study the relations between putative DMN subnetworks and attentional behaviour.

Stereoelectroencephalography (sEEG) utilizing stereotactically implanted intracranial electrodes provides a unique window to study moment-to-moment attentional activity. These electrodes are commonly implanted for delineation of seizure-initiating brain regions in a clinical population of children with epilepsy, who are predisposed towards attentional deficits (McKhann, 2020; Youngerman et al., 2019). Local-field potentials (LFPs) may be directly recorded from across the brain including difficult to access regions known to subserve key roles in attentional behaviour, including insula, ACC/PCC, and mPFC (Youngerman et al., 2019). In comparison to non-invasive methods, these intracranial recordings permit granular examination of processing within and between key attentional nodes at relevant neural timescales and afford significantly greater spatial precision with which to examine putative functional specialization within large-scale neural networks (Voloh et al., 2015; Womelsdorf et al., 2007; Wong et al., 2021). However, studies utilizing sEEG to study attentional behaviour have been limited due to significant inter-subject heterogeneity in implantations strategies, which are dictated by clinical epilepsy monitoring requirements.

The examination of subject-specific neural activation patterns represents one approach with which to resolve such cross-implantation heterogeneity. Deep learning is an artificial intelligence tool that may be applied toward sEEG data to identify such subject-specific neural patterns without the need to program pre-exiting assumptions about relevant features. Preliminary data suggest that a single-subject approach may also reveal a more detailed picture of moment-to-moment neural interactions lost in aggregate analyses (Cohen and Cavanagh, 2011; Pandarinath et al., 2018; Vahid et al., 2020). To date, applications of deep learning modelling within neuroscience have been limited largely due to an inability to delineate the features utilized to generate model predictions. However, advanced model-agnostic algorithms now permit detailed study of neural correlates associated with model predictions, significantly increasing the value of deep learning applications within the study of attention (Lundberg and Lee, 2017).

Given converging evidence for (i) moment-to-moment modulation of cognitive flexibility within key DMN subsystems, (ii) the ability to robustly sample this circuitry through sEEG and (iii) the potential for deep learning to reveal common patterns of neural activation, despite considerable inter-subject variability within datasets, we sought to characterize the network substrates of attentional set shifting by applying deep learning to intracranial recordings. Using a custom convolutional neural network (CNN), individualized predictive models based on modulations in single-trial power were generated and characterized through large-scale network mapping. The results of this work provide novel insights into the dissociable network subsystems that subserve attentional behaviour in children. We also provide a novel analytic framework enabled by deep learning to study neurocognitive processes in heterogenous datasets of intracranial recordings.

## 2. Results

### 2.1 Unique sEEG Cohort

Data were obtained from 608 unique recording sites across a consecutive series of thirteen children admitted for invasive epilepsy monitoring at the Hospital for Sick Children Epilepsy Monitoring Unit (EMU). To ensure that our results could not likely be attributed to epileptiform activity, recordings were performed from outside epileptogenic zones and at least 12 hours from clinical or electrographic seizure events. Trials coincident with interictal discharges and significant artifacts were excluded from further analysis (see Methods).

Attentional performance was indexed during an attentional set-shifting task extensively validated in paediatric cohorts (Figure 1A) (Oh et al., 2014). There are two trial conditions in this paradigm: i) shift trials, in which response rules change implicitly from prior trials, and ii) non-shift trials, in which response rules remain unchanged. During the shift condition, selective attention and mental flexibility are essential to adapt responses accurately and efficiently (Oh et al., 2014). The non-shift condition does not engage cognitive flexibility and instead measures sustained attention (Oh et al., 2014). In both trial types, slowed reaction times (RT) and errors indicate lapses in attention. Comparison of these distinct trial types therefore permits selective examination of neural responses associated with mental flexibility during attentional shifts (Oh et al., 2014).

**Figure 1.**
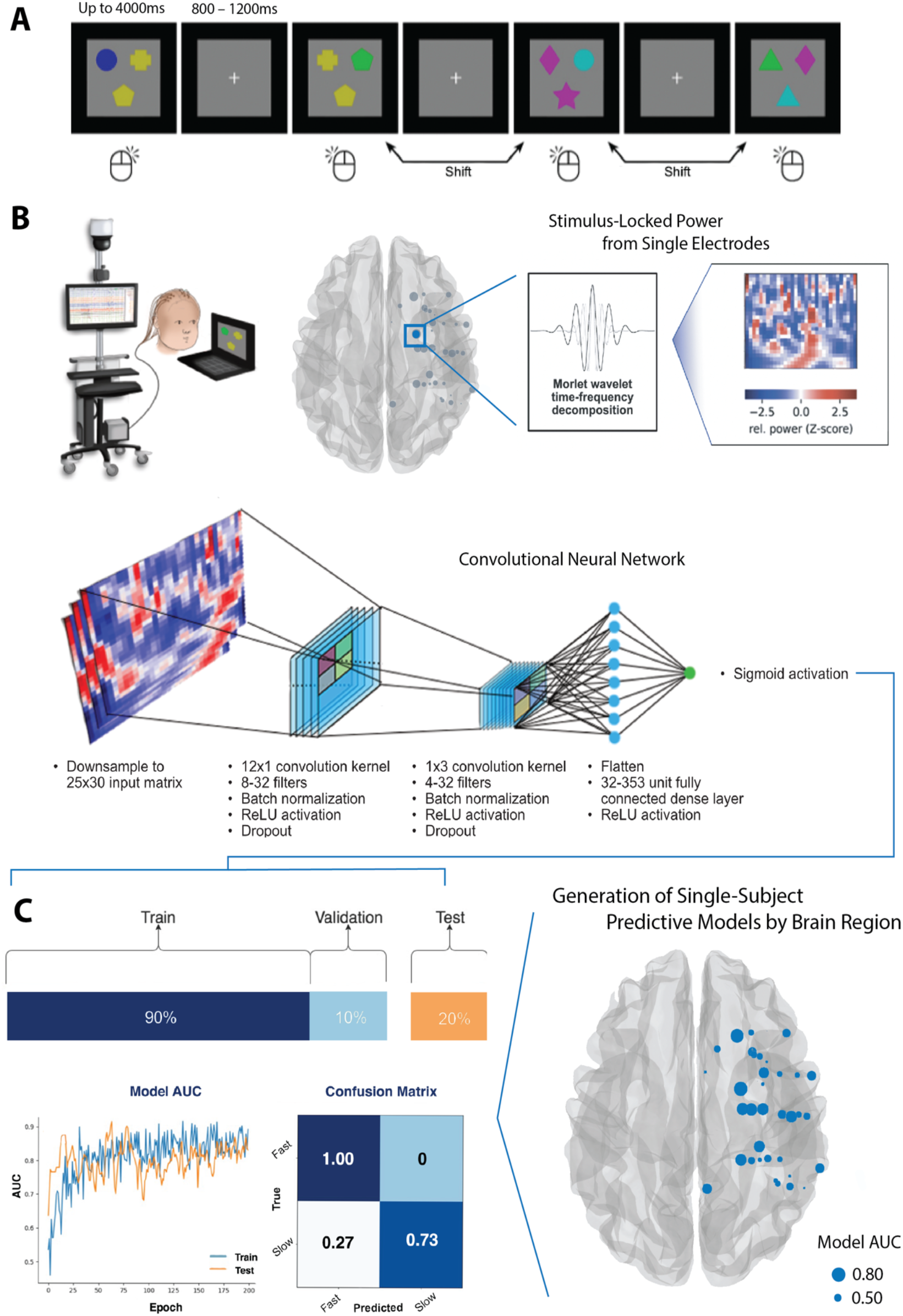
Overview of set-shifting task, data acquisition, and deep learning architecture. **A**. Schematic overview of the computer-based attentional set-shifting task (Oh et al., 2014). **B**. Intracranial EEG data from all implanted electrodes is recorded during task performance. In order to delineate brain regions predictive of attention, separate models are then trained on individual electrode data within subjects. The input features for the classifier are stimulus-locked spectral power obtained by Morlet continuous wavelet transform. **C**. Each brain-region specific model is tuned using 10-fold cross-validation and attentional performance scored using a held-away test set. The predictive ability of each individual brain region is then indexed by model AUC score and can be visualized as shown on the right of the panel, where node size equates to model AUC score.

### 2.2 Beta and gamma power in key attentional networks predict trial performance

Deep learning models were trained on single-trial sEEG data from individual brain regions within each subject (see Methods for data processing approach). These models demonstrate that modulations in stimulus-locked power could accurately predict attentional performance as indexed by RT. Given the individualized approach employed, specific brain regions and power features most predictive of performance varied between subjects. In order to determine the neuroanatomical substrates of attentional behaviour by trial type, performance of each individual classifier was aggregated across all subjects and grouped by the functional network/brain region sampled. Only brain regions with aggregate AUC values significantly greater than chance-level were included for further analysis (q < 0.05 with FDR correction, one-sample t-test).

The application of Shapley explanations (Lundberg and Lee, 2017) to these models permitted detailed analysis of the neural features learned by each classifier (Figure 2A; see Methods for detailed explanation). Across brain regions, modulations in single-trial beta power were the greatest contributor to model predictions explaining 47% of prediction variance across both shift and non-shift trials (Tables S1 and S2). Gamma-power was the next major contribution (25%), followed by alpha-(19%) and theta-band activity (9%).

**Figure 2.**
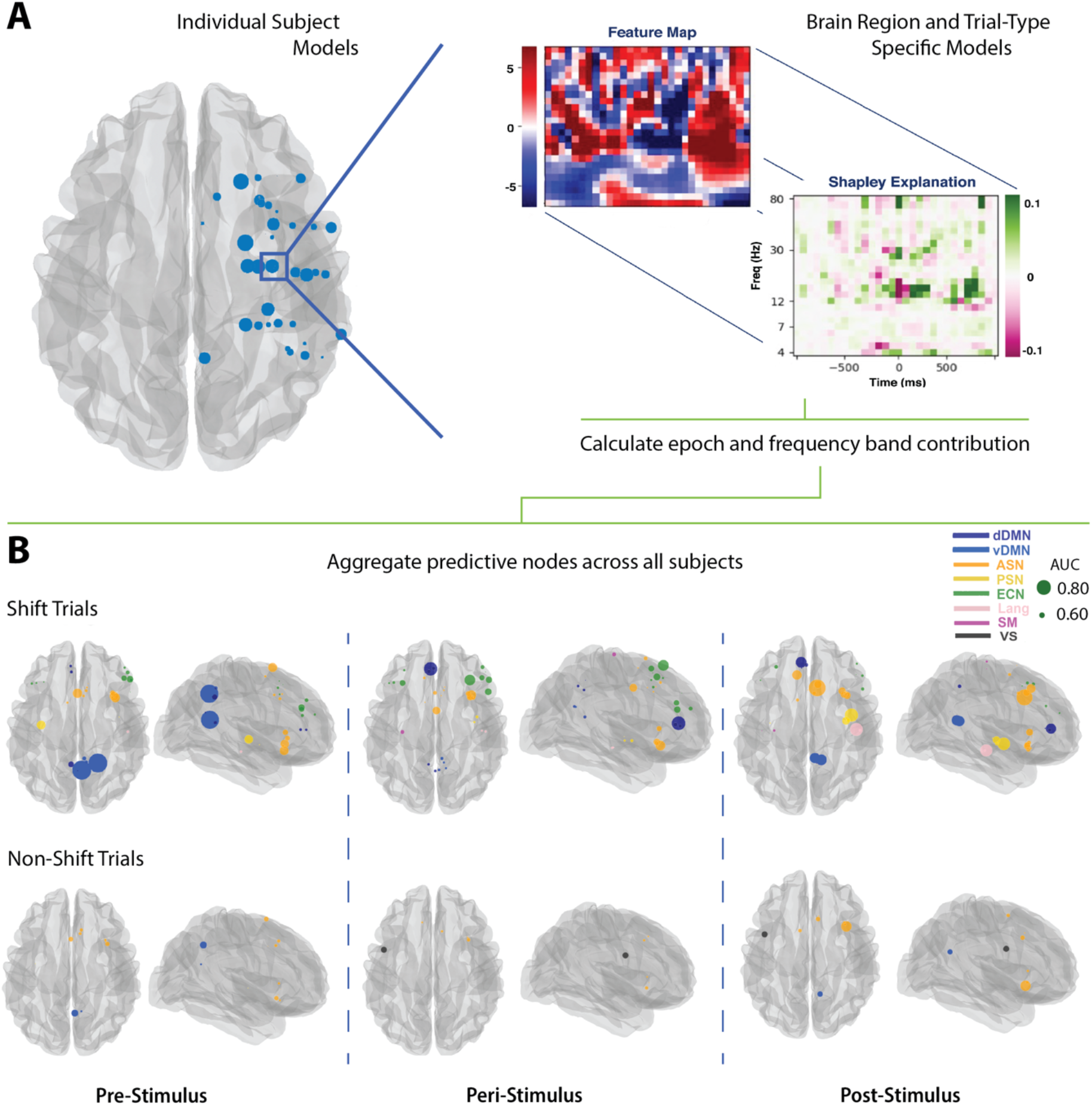
Overview of model explanation approach and summary of predictive nodes for shift and non-shift trials. **A**. For each node, Shapley value-based feature importance was calculated highlighting features that supported the prediction of slow RT (green) and those that argued toward a prediction of fast RT (magenta). Within each model, the contribution of power within defined frequency bands and time windows was calculated. **B**. Aggregate data in the beta-band across all patients are shown. Across patients, only those regions with AUC significantly above chance are shown (one-sample t-test, comparator = 0.5). During attentional shift trials, modulations in beta power within the dorsal DMN, ECN, and SN were predictive of attentional performance (q < 0.05, FDR-corrected). In trials without attentional shift, a smaller number of nodes within the ventral DMN, SN and visuospatial networks were predictive of RT (q < 0.05, FDR-corrected). <u>Abbreviations</u>: dDMN/dorsal DMN; vDMN/ventral DMN; ASN/Anterior SN; PSN/posterior SN; Lang/language network; SM/somatomotor network; VS/visuospatial network.

During attentional shift trials, beta- and gamma-band power within nodes of the ECN (inferior/middle frontal), SN (insula, middle cingulate, and supplementary motor area), and DMN (ACC, precuneus) predicted performance (Table S1; q < 0.05). Figure 2B demonstrates the spatial distribution and performance of predictive models based on modulations in beta-band power. Fewer brain regions were predictive of RT in non-shift trials. In this condition, only activity within nodes of the SN (insula, supplementary motor area) and ventral DMN (precuneus) predicted performance (Table S2; q < 0.05). Uniformly, predictive models for the attentional shift condition performed better than those for trials that did not require shifts (AUC_shift_: 0.74 ± 0.01; AUC_non-shift_: 0.62 ± 0.01 – mean AUC of all nodes ± std. error; p < 0.001 independent samples t-test).

### 2.3 Shifts in attention preferentially engage dorsal DMN

We subsequently sought to analyze the functional networks engaged by highly predictive attentional nodes during task performance using connectomic profiling. Briefly, a 3mm region-of-interest was created about the MNI coordinate of each highly predictive node. As previously described, whole-brain connectivity maps for each sphere were then derived using a publicly available 3-Tesla normative resting-state fMRI dataset (Thomas Yeo et al., 2011). Statistical connectivity maps describing voxel-wise correlations with each seed region were derived from low-frequency BOLD fluctuations across the 1,000 subjects included in this dataset and corrected for multiple comparisons with whole-brain Bonferroni correction at pBonferroni < 0.05 (Elias et al., 2020; Joutsa et al., 2018; Mithani et al., 2019). As a significant degree of overlap existed in the neural circuitry engaged by the shift and non-shift trial conditions, voxelwise odds ratios (VORs) were generated to analyze functional connectivity patterns preferentially associated with the attentional shift vs. non-shift conditions (see Methods for further detail). Network engagement was examined during three processing epochs relative to stimulus onset (pre-stimulus: −1,000ms to −250ms; peri-stimulus: −250ms to 250ms; and post-stimulus: 250 to 1,000ms), due to prior evidence supporting critical neural interactions during these distinct time windows (Wong et al., 2021). Schematic overview of this analysis pipeline is shown in Figure 3A.

**Figure 3:**
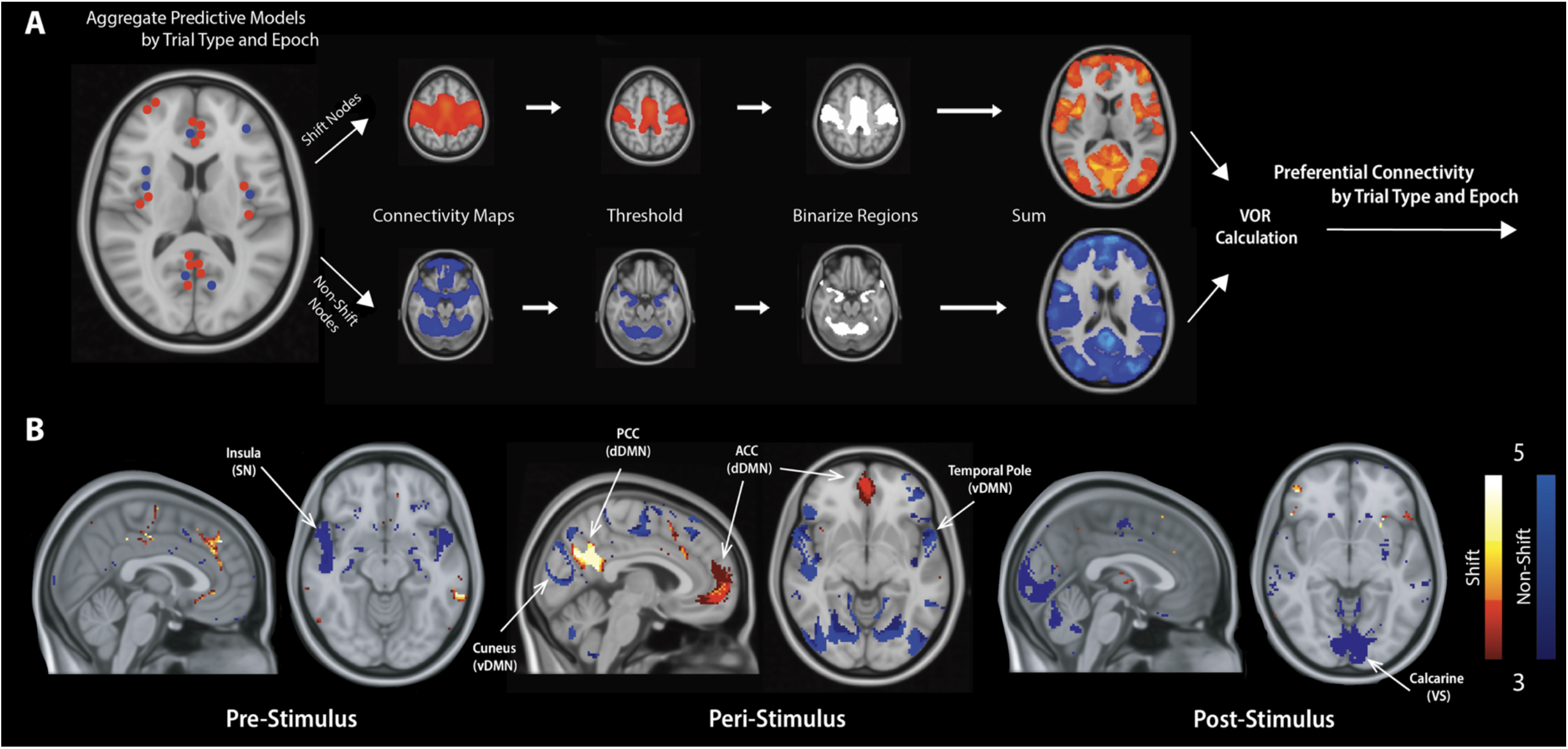
Voxelwise odds-ratio maps of functional networks engaged during task performance. **A**. Highly predictive nodes were aggregated across subjects. Whole-brain connectivity patterns were then determined in relation to these nodes using a publicly available resting-state fMRI dataset (http://neuroinformatics.harvard.edu/gsp/) (Thomas Yeo et al., 2011). A conservative threshold at pBonferroni < 0.05 was then applied and connectivity maps were binarized. VORs for the shift and non-shift conditions were then calculated as described in Eqs. 1 and 2. **B**. During attentional shift trials, predictive nodes preferentially engage ACC and PCC within the dorsal DMN during stimulus processing (mean VOR = 2.04). During the same epoch, trials without attentional shift demonstrate preferential engagement of ventral DMN (mean VOR = 2.20). In non-shift trials, highly predictive nodes also engage the insula (mean VOR = 2.39/R; 2.13/L) in the pre-stimulus period and calcarine cortex (mean VOR = 2.38/R; 2.41/L) in the post-stimulus period. <u>Abbreviations</u>: dDMN/dorsal DMN; vDMN/ventral DMN; VS/visuospatial network.

Brain regions predictive of RT in attentional shift trials showed robust engagement of the dorsal DMN during task performance. During the peri-stimulus epoch, dorsal DMN engagement was more than twice as likely during shift trials in comparison to those without an attentional shift (Figure 3B; mean VOR = 2.04, max VOR = 5.25). Within this network, right ACC (mean VOR = 2.01) and right PCC (mean VOR: = 2.77) were preferentially engaged during stimulus processing (Figure 3B). No other large-scale network engagement patterns were observed during attentional shift trials.

In contrast with engagement of the dorsal DMN engagement during shift trials, non-shift trials engaged the ventral DMN during peri-stimulus periods (Figure 3B; mean VOR = 2.20, max VOR = 23.4). Within ventral DMN, a variety of brain regions were preferentially engaged, including right amygdala (mean VOR = 3.27), bilateral cuneus (mean VOR = 3.05/R; 3.71/L), temporal pole (mean VOR = 6.31/R; 4.36/L), and left inferior temporal gyrus (mean VOR = 4.06). In non-shift trials, pre-stimulus insular engagement (mean VOR = 2.39/R; 2.13/L) and post-stimulus calcarine engagement were also observed (mean VOR = 2.38/R; 2.41/L). Across both trial conditions, the dorsal/ventral DMN were only engaged during the peri-stimulus period (Figure 3B).

## 3. Discussion

The mechanisms subserving cognitive flexibility and shifts in attention remain poorly understood. Impairments in mental flexibility have been reported across the spectrum of attentional disorders (Dajani and Uddin, 2015; Genet and Siemer, 2011; Luna-Rodriguez et al., 2018; Roshani et al., 2020). In children, higher cognitive flexibility scores correlate with improved quality of life as well as future academic, social, and professional accomplishment (Dajani and Uddin, 2015). Here we apply deep learning to intracranial recordings during an attentional set-shifting task to delineate modulations in stimulus-locked power associated with mental flexibility at the single trial and single subject level. Based on these models, we further examine the spatial and temporal patterns of large-scale networks engaged by highly predictive attentional nodes.

This approach yielded three main findings relevant to the study of human attention. First, we show that trial-by-trial beta and gamma modulations within ECN, SN, and DMN are predictive of subject performance in a set-shifting task. Furthermore, we demonstrate that neural centres critical for attentional shift performance selectively engage the dorsal DMN. Finally, we demonstrate that subject-specific deep learning algorithms trained on single-trial stereotactic LFP data can learn biologically relevant parameters and converge upon consistent large-scale functional networks despite inter-subject variability in implantation. Taken together, these findings implicate the dorsal DMN stream in childhood cognitive flexibility and provide novel machine learning approaches to study granular attentional circuitry.

The existence of functionally distinct streams within the DMN provides a parsimonious hypothesis for the role of this critical network in cognitive flexibility and attention. Classical conceptualizations of the DMN as a task-negative network primarily involved in self-referential and introspective processing (Buckner et al., 2008; Davey et al., 2016; Kucyi and Davis, 2014; Raichle, 2015) have been challenging to reconcile with more recent reports of network recruitment during externally-directed attention (Crittenden et al., 2015; Smith et al., 2019, 2018; Waltz et al., 2013). Contextual encoding models offer a potential explanation, proposing that the DMN responds to degrees of change in cognitive context (Crittenden et al., 2015; Smith et al., 2018). In this contextual framework, DMN engagement reflects “relaxation” of attentional foci to facilitate mental shifts. Network activation at rest, particularly during mind-wandering, would be explained by ongoing shifts in thought processes and their mental context (Kucyi and Davis, 2014; Mittner et al., 2016; Zhou and Lei, 2018). The existence of distinct subsystems within human DMN provides strong support for functional specialization within this network that may dually subserve contextual encoding (dorsal stream) and introspective processing (ventral stream).

The deep learning models consistently identified beta and gamma power as reliable predictors of attentional performance (Tables S1/S2). In non-human primates, dorsal DMN (ACC) has previously been shown to facilitate mental flexibility through coupling of low-frequency phase with both beta- and gamma-band power distributed across attentional nodes (Voloh et al., 2015; Womelsdorf and Everling, 2015). Recent data suggest that a similar mechanism may also be implicated in human attention (Wong et al., 2021). The identification of these specific frequency bands in the absence of prespecified feature selection strongly supports a role for neocortical beta and gamma oscillations in distributed attentional control (Fries, 2015; Hanslmayr et al., 2009; Kopell et al., 2000; Sederberg et al., 2006; Spitzer and Haegens, 2017). In ECN and dorsal DMN, beta and gamma power were specifically associated with attentional shifting while insular power (SN) predicted performance regardless of trial type. Such a result is consistent with lower-order stimulus processing within the SN (Chi et al., 2019; Jiang et al., 2000; Lenartowicz and Loo, 2014; Szuromi et al., 2011). During attentional shifts, distributed beta/gamma activity coordinated by ECN and DMN hubs may serve to facilitate shifts in focus.

Recent work by Vu *et al*. (Vu et al., 2018) has recognized the potential of deep learning in neuroscience while acknowledging the need for increased model interpretability. In the current work, we demonstrate the application of a model-agnostic approach based on game theoretic Shapley values (Lundberg and Lee, 2017). The use of Shapely values allowed us to determine the gain of individual features toward final model prediction. These model explanations were then cross-referenced with trial-by-trial time frequency decompositions to delineate the features learned by each classifier (Figure 2A). A biologically relevant patten in power modulations across attentional nodes was identified without prior explicit programming. Furthermore, the utilization of deep learning allowed us to develop patient-specific attentional models that could identify individual-specific neural rhythms and resolve cross-montage heterogeneity. As the study of cognition using intracranial recordings grows, targeted applications of deep learning models combined with model-agnostic feature explanation algorithms hold significant utility toward revealing the neural syntax associated with specific behavioural states.

The present study utilizes a combined approach leveraging deep learning and sEEG to study the dynamic neural syntax associated with human attention and cognitive flexibility. First, we demonstrate that single-trial power within nodes of the ECN, SN, and dorsal DMN predicts performance during shifts in attention. We further demonstrate that attentional nodes engage dorsal DMN hubs within the ACC/PCC in association with mental flexibility; ventral DMN hubs in precuneus and temporal lobe are engaged during default responses. Finally, we report a method for co-application of deep learning and intracranial electrode recording that can be leveraged to overcome challenges associated with between-subject heterogeneity inherent to human intracranial data. The existence of distinct functional streams within the DMN provides a parsimonious account for both the contextual encoding and default-mode responses observed within this critical network. These data further provide an interpretable deep learning framework for the study of human attention and associated cognitive domains.

## 4. Materials & methods

### 4.1 Subjects

Thirteen consecutive children admitted for invasive epilepsy monitoring at the Hospital for Sick Children Epilepsy Monitoring Unit (EMU) were included in the study. Clinical and demographic data for all included subjects are provided in Table 1. Subjects were prospectively recruited into the study and study procedure was fully approved by an independent ethics review board at the Hospital for Sick Children. Informed consent and assent were obtained from all patients and their caregivers in compliance with the Code of Ethics of the World Medical Association (Declaration of Helsinki). All patients were diagnosed with medically intractable epilepsy and underwent surgical placement of sEEG depth electrodes (0.86mm diameter; AdTech, USA) for clinical seizure mapping. The location, laterality, and number of electrodes implanted was dictated solely by clinical needs and varied between patients.

**Table 1:**
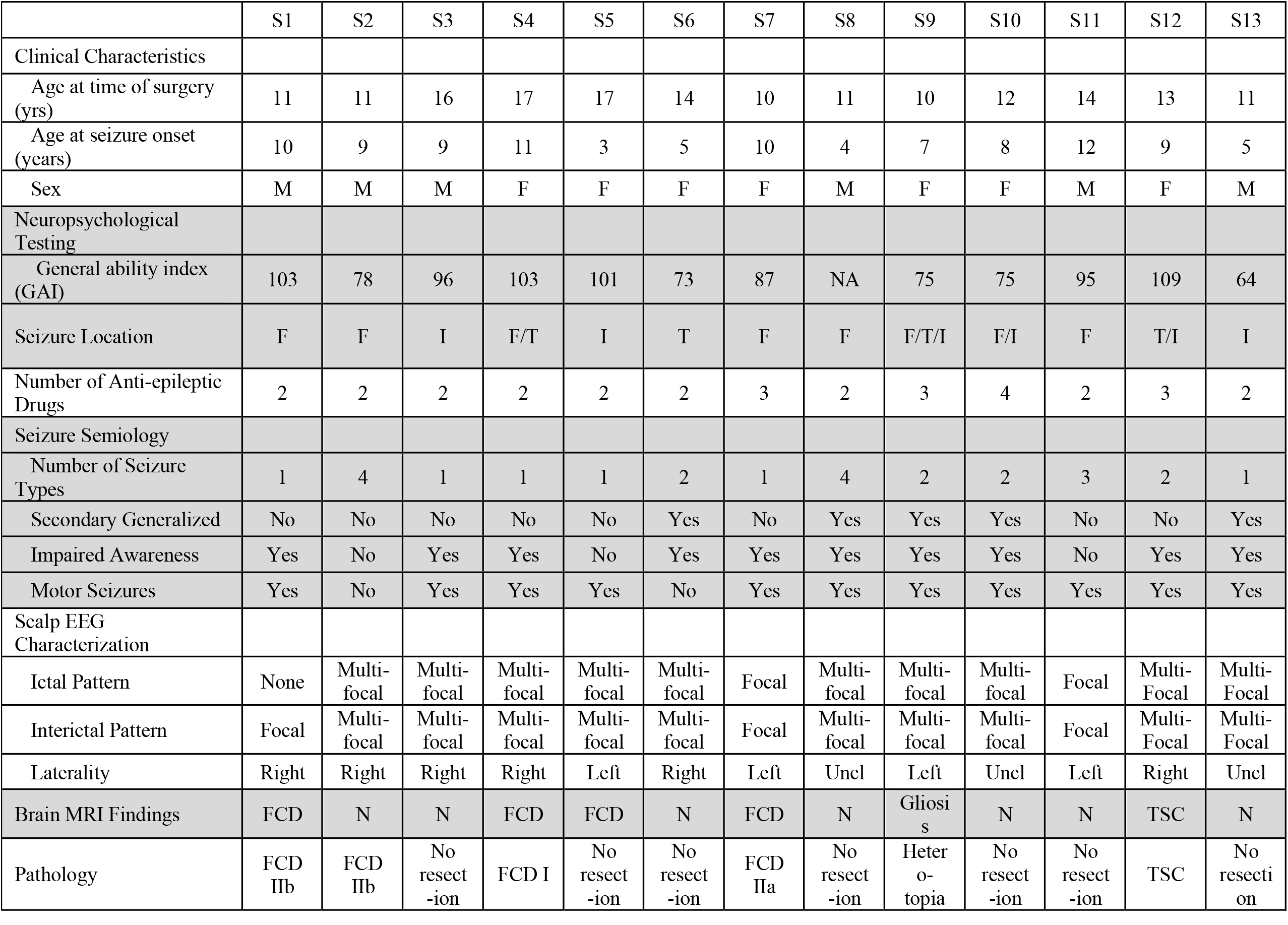
Clinical and demographic data for the included patients. F-frontal; FCD-focal cortical dysplasia; I-insular; N-normal; T-temporal; TSC – tuberous sclerosis complex; Uncl - unclear

### 4.2 Intracranial electrode localization and image processing

The intracranial electrodes were localized and transformed into standard Montreal Neurological Institute (MNI) space using a published protocol (Stolk et al., 2018). In brief, the pre-operative T1 3-Tesla magnetic resonance imaging (MRI) and post-operative computed tomography (CT) scans of each patient were aligned to the AC-PC (anterior commissure, posterior commissure) coordinate system and linearly co-registered. The electrode locations were visually marked on the co-registered CT scan using the Fieldtrip toolbox. Finally, the 3D coordinates of the electrode contacts were non-linearly warped onto a template brain in MNI space. Each electrode contact was labelled based on the anatomic brain and region the resting-state functional network sampled using the AAL (Rolls et al., 2020) and FIND (Shirer et al., 2012) atlases respectively.

### 4.3 Attentional testing

Attentional performance was indexed during an attentional set-shifting task (Oh et al., 2014). Given the body of literature linking set-shifting impairments to childhood-onset attention deficits (Kalkut et al., 2009; Luna-Rodriguez et al., 2018; Oades and Christiansen, 2008), this task has been validated for the study of attention in children. A schematic overview of the task is shown in Figure 1A. During the task, participants are required to match one of two coloured images with a third stimulus (target) according to either its colour or shape. The task consists of two trial conditions: i) “non-shift” trials during which the colour or shape of the target remains constant, and ii) “shift” trials in which the color or shape rule changes, requiring an attentional shift. Set-shifting is a complex measure of sustained attention and cognitive flexibility that relies on the complimentary processes of attentional shift, response inhibition, and salience detection (Dajani and Uddin, 2015; Luna-Rodriguez et al., 2018; Periáñez et al., 2004). In this task paradigm, selective attention and mental flexibility are essential to accurately and efficiently adapt responses during shift trials. Slowed reaction times and errors indicate lapses in attention. Trials ended when a button press was registered or after 4,000ms. The interstimulus interval between trials was jittered between 800-1,200ms (mean of 1,000ms)

All subjects were trained to accurately perform the task and trial by trial fluctuations in attentional performance were indexed by RT. The trial-wise RT was chosen as the primary endpoint, while accuracy was maintained. This was performed given that variability in RT is a more reliable determinant of attention than other measures such as task accuracy or error rates (Kalff et al., 2005). Furthermore, functional limitations in domains such as academic skills, adaptive behaviour, and social skills are strongly correlated with RT variability in children with attentional disorders (Cook et al., 2018).

### 4.4 EEG preprocessing

All subjects underwent digital recording of sEEG signals using Natus Quantum 256-channel amplifier (Natus Medical Inc., Pleasanton, CA, USA) with a sampling rate of 2048 Hz. The data were exported into Python-compatible formats and all signals processing was performed using MNE-python package (Gramfort et al., 2013). A notch-filter was applied to remove line noise and harmonics and the data were re-referenced to a common average reference montage. EEG data for individual trials were temporally locked to target presentation. Trimmed segments of the EEG spanning from one second pre-to one second post-stimulus were used to develop the input features to our machine learning classifier. The final data were visually examined to ensure artefact removal.

To control for the effects of epileptiform events, all recordings were performed at least 12 hours from a clinical or electrographic seizure event. Trials containing epileptiform interictal discharges and significant artifacts were removed from the analysis (on average 0-5% of total trials per patient) based on automated detection algorithms (Jas et al., 2017; Wong et al., 2021).

### 4.5 Input features

The input features for the CNN were single-trial time-frequency decompositions of spectral power time-locked to stimulus onset as described above. Five-cycle Morlet continuous wavelet transform over 25 logarithmically spaced centre frequencies ranging from 4-80Hz was used to compute spectral time-frequency decompositions (Gramfort et al., 2013). To reduce the dimensionality of the feature space, the signal was downsampled in the temporal domain using a 60ms sliding window with an overlap of 50%. Finally, as deep learning approaches are sensitive to scaling, the power values were z-scored relative to the mean and standard deviation in power of each frequency across time. At a single trial level, the resulting feature matrix for the classifier thus represented the z-scored spectral power at 25 frequencies across 30 timepoints. An example of the feature inputs to the model is shown in Figure 1B.

### 4.6 Deep learning classifier

In the first phase of this analysis, a custom CNN was built using the Tensorflow/Keras framework in Python (Figure 1B) (Chollet, 2015). The goal of the classifier was to predict a patient’s attentional performance as indexed by a “fast” or “slow” RT on the given trial. The dichotomization between “fast” and “slow” trials was based on the median reaction time for each patient to maintain balanced strata. Furthermore, trials with attentional shift were separated from those without attentional shift and different models were trained for each of these conditions.

For each patient, deep learning classifiers were trained based on data from individual intracranial contacts representing different brain regions. This single-contact approach allowed us to determine the contribution of a specific brain region to each subject’s attentional performance. Ten-fold cross-validation was utilized to select the optimum hyperparameters for each model. The final performance of each brain region-specific classifier was evaluated on an independent held-away test set comprising 20% of the data. Figure 1C demonstrates typical training curves for a model, the data proportioning approach used, and a spatial representation of all brain region-specific models trained for a single patient. The performance of each region-specific attentional model was scored based on accuracy and area under the receiver operating characteristic curve (AUC) metrics. To examine whether modulations in single-trial power were predictive of attention, data from individual brain-region specific classifiers were aggregated. Across subjects, AUC metrics for all models sampling the given brain region were pooled and compared to chance.

### 4.7 Model-agnostic feature explanation

Using SHAP package in Python (Lundberg and Lee, 2017), we calculated the feature importance for spectral power at each time and frequency point, as shown in Figure 2A. Prediction contribution in the theta-(4-8Hz), alpha-(8-12Hz), beta-(12-30Hz), and gamma-bands (30-80Hz) was determined. Feature importance was also epoched based on stimulus processing windows previously implicated in human set-shifting: i) pre-stimulus (−1,000 to −250ms), ii) peri-stimulus (−250 to 250ms), and iii) post-stimulus (250 to 1,000ms) (Wong et al., 2021). The relative weight of model contributions within each time window and frequency band was then calculated, adjusted for window length. Time-window weighted model data were used in the following step to compute time-window specific engagement of relevant functional networks.

### 4.8 Connectomic profiling

Subject-specific deep learning models provided a set of highly predictive electrode nodes that were related to performance on the attentional set shifting task. To better understands the normative neural networks engaged by these disparate nodes, a connectomic analysis was performed on all electrode notes that yielded a predictive value of AUC > 0.7. A 3mm region-of-interest was created about the coordinate of each highly predictive electrode node in MNI space. We then derived whole brain connectivity maps for each sphere using a publicly available high-quality 3-Tesla normative resting-state fMRI dataset (http://neuroinformatics.harvard.edu/gsp/) (Thomas Yeo et al., 2011). As previously described (Elias et al., 2020; Joutsa et al., 2018; Mithani et al., 2019), connectivity r-maps describing the voxel-wise correlations with each seed region were derived from the time course of low-frequency BOLD fluctuations across the 1,000 subjects included in this dataset (healthy controls; 57% female, age range: 18-35y) using an in-house MATLAB script (The MathWorks, Inc., Version R2019b. Natick, MA, USA). The r-maps were then converted to t-maps and corrected for multiple comparisons (Bonferroni correction voxel-wise for the whole-brain) using the known p-value distribution at t=5.1 (pBonferroni < 0.05). These thresholded maps were then binarized preserving the significant brain-network and topographical relationship associated with each seed (pBonferroni < 0.05). This conservative strategy was applied to overcome limitations associated with applying normative functional connectomic data to epilepsy patients, whose functional connectivity may significantly deviate from healthy controls (Ibrahim et al., 2014; Pittau et al., 2012). Stringent statistical thresholding and binarization aims to remove the influence of subtle connectivity patterns present in normative connectomic data less likely to hold true to epilepsy patients (Mansouri et al., 2020; Pittau et al., 2012). An overview of this pipeline is shown in Figure 3A.

To investigate patterns of network engagement preferentially associated with attentional shift, VOR maps were constructed from the binarized brain-wide functional maps associated with each seed. The VOR maps were calculated as previously described (Boutet et al., 2018; Sprenger et al., 2012). Equation 1 shows the formula used to calculate the shift vs. non-shift VORs, while equation 2 shows the non-shift vs. shift VOR calculation.

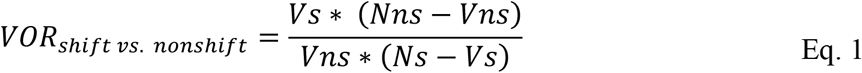

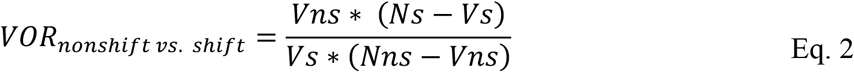

In the above equations, Ns = number of seeds in shift condition p; Nns = number of seeds of non-shift condition c; Vs = number of seeds in shift trials overlapping a specific voxel; Vns = number of seeds in non-shift trials overlapping a specific voxel. The VOR values as calculated above give the likelihood of a voxel to be preferentially engaged during shift or non-shift trials respectively. Therefore, a VOR value of 2.0 for the shift vs. non-shift comparison indicates that a given voxel is twice as likely to be involved in shift vs. non-shift trial conditions.

## Supplementary Material

### Mean reaction times

The distribution of reaction times for the 13 subjects included in the analysis are presented in **Figure S1.**

**Figure S1:**
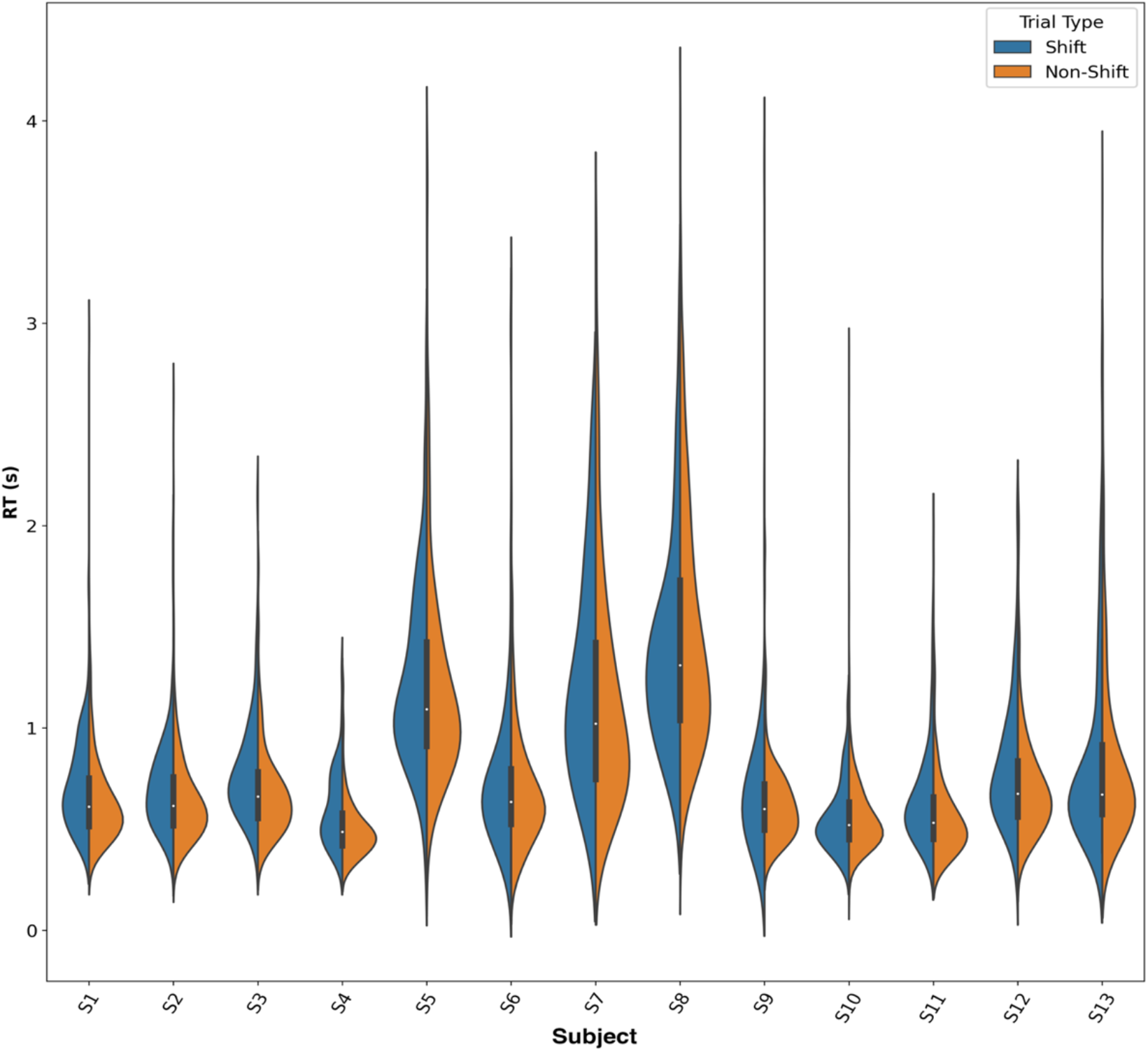
Distribution of reaction times per subject. Violin plots demonstrate per-subject reaction times on the attentional set-shifting task. The distribution of reaction times for the shift and non-shift attentional conditions is shown in blue and orange, respectively. Note that the distribution of reaction times for the task varies considerably between subjects and trial types.

### Single-trial modulations in power predict attentional performance

To determine the features predictive of attention, data from each individualized and brain-region specific classifier was examined. The anatomic brain region and functional network parcellation of the node utilized by each model was determined using the AAL and FIND atlases as described. We then aggregated these models across all subjects and grouped them together based on the functional network/brain region sampled. To ensure that our analysis focused only on regions with significant predictive ability across subjects, only brain regions with aggregate AUC values significantly greater than chance-level (0.50) were included for further analysis (q < 0.05 with FDR correction, one-sample t-test). Finally, using the described model explanation algorithms, the relative contributions of power within the theta, alpha, beta, and gamma bands was calculated and averaged across models sampling the given brain region/functional network pair. These data are presented in **Tables S1/S2**.

**Table S1.** Overview of functional networks and brain regions predictive of attentional shift. Only regions with an aggregate AUC significantly greater than chance level are shown (q < 0.05, FDR corrected). Theta, alpha, beta, and gamma columns indicate the proportion of model prediction explained by nodulations in single-trial power in the given frequency band. Bold indicates the frequency band of greatest contribution.

|  |  | AUC | Theta | Alpha | Beta | Gamma |
| --- | --- | --- | --- | --- | --- | --- |
| **FIND Network** | **Region** |  |  |  |  |  |
| Left ECN | Frontal_Mid_L | 0.77 | **0.47** | 0.22 | 0.18 | 0.14 |
| Language | Temporal_Mid_R | 0.80 | 0.35 | 0.14 | **0.46** | 0.05 |
| Right ECN | Frontal_Inf_Tri_R | 0.68 | 0.03 | 0.19 | **0.52** | 0.26 |
| Right ECN | Frontal_Mid_R | 0.66 | 0.07 | 0.11 | **0.45** | 0.38 |
| Sensorimotor | Precentral_L | 0.75 | 0.04 | 0.00 | 0.16 | **0.79** |
| Ant. SN | Cingulum_Mid_R | 0.78 | 0.19 | 0.21 | **0.43** | 0.17 |
| Ant. SN | Insula_R | 0.71 | 0.04 | 0.23 | **0.57** | 0.16 |
| Ant. SN | Supp_Motor_Area_L | 0.75 | 0.05 | 0.10 | **0.50** | 0.35 |
| Dorsal DMN | Cingulum_Ant_L | 0.78 | 0.02 | 0.26 | **0.48** | 0.23 |
| Dorsal DMN | Precuneus_L | 0.72 | 0.06 | 0.14 | 0.30 | **0.49** |
| Post. SN | Insula_L | 0.68 | 0.00 | 0.29 | **0.50** | 0.21 |
| Post. SN | Insula_R | 0.71 | 0.06 | 0.16 | **0.50** | 0.28 |
| Ventral DMN | Precuneus_R | 0.79 | 0.12 | 0.16 | **0.45** | 0.28 |

**Table S2.** Overview of functional networks and brain regions predictive of attention in non-shift trials. Only regions with an aggregate AUC significantly greater than chance level are shown (p < 0.05, FDR corrected).

|  |  | AUC | Theta | Alpha | Beta | Gamma |
| --- | --- | --- | --- | --- | --- | --- |
| **FIND Network** | **Region** |  |  |  |  |  |
| Visuospatial | Precentral_L | 0.62 | 0.05 | 0.26 | **0.42** | 0.27 |
| Ant. SN | Cingulum_Mid_R | 0.65 | 0.05 | 0.21 | **0.70** | 0.05 |
| Ant. SN | Insula_R | 0.61 | 0.04 | 0.31 | **0.49** | 0.16 |
| Ant. SN | Supp_Motor_Area_L | 0.64 | 0.23 | 0.19 | **0.41** | 0.17 |
| Post. SN | Insula_L | 0.57 | 0.06 | 0.02 | **0.47** | 0.45 |
| Ventral DMN | Precuneus_R | 0.61 | 0.13 | 0.14 | **0.62** | 0.11 |

## Notes

### Competing Interest Statement

The authors have declared no competing interest.

